# Chromosome-level genome assembly of *Calotes wangi* with dynamic colour variation

**DOI:** 10.64898/2026.07.07.736949

**Authors:** Xia Qiu, Yiru Wang, Jieru Wen, Ying Chen, Longhui Zhao, Jiayi Jian, Weizhao Yang

## Abstract

The Wang’s garden lizard, *Calotes wangi*, is a widely distributed agamid species in Southern China and Northern Vietnam and exhibits pronounced colour variation and rapid body colour change. Despite increasing interest in the genomic basis of colour variation, chromosome-level genomic resources remain limited in agamid lizards. Here, we generated a chromosome-level reference genome of *C. wangi* using PacBio HiFi sequencing and Hi-C scaffolding. The final genome assembly was approximately 1.66 Gb in size and comprised 6 macrochromosomes and 11 microchromosomes, with a contig N50 of 110.09 Mb and 98.9% complete BUSCO genes. A total of 20,442 protein-coding genes were annotated. Comparative genomic analyses identified 297 significantly expanded gene families, with enriched functions associated with steroid metabolism, chromatin regulation, and epigenetic processes. This high-quality genome assembly provides an important genomic resource for future studies of colour variation, phenotypic plasticity, and evolutionary diversification in agamid lizards.

## Background & Summary

Animal colouration plays important roles in camouflage, thermoregulation, and social communication^1,2^. In many species, colouration also functions as an important visual signal during territorial interactions and mate choice^3–5^. Recent chromosome-level genomic studies have demonstrated that structural genomic variation and regulatory evolution contribute significantly to phenotypic and colour diversification in animals, including camouflage adaptation in stick insects and reproductive morph variation in ruffs^6,7^. In contrast, comparable chromosome-level genomic resources remain limited in lizards with diverse colouration patterns, particularly within the family Agamidae^8–10^.

The Wang’s garden lizard, *Calotes wangi* Huang et al., 2023^11^, is a widespread agamid species distributed in Southern China and Northern Vietnam^11–14^. Recent taxonomic and phylogenetic studies have indicated that *C. wangi* belongs to the *C. versicolor* species complex and is genetically distinct from other members of the complex^11,12^. *C. wangi* exhibits pronounced context-dependent colour variation and rapid body colour change^15^. Individuals can shift between green and brown colour morphs under different environmental conditions. Males often develop conspicuous reddish or orange colouration during social interactions and the breeding season^15^. Its pronounced colour plasticity and phenotypic variability make *C. wangi* a valuable system for studying the genomic basis of colour variation and evolutionary diversification.

To date, only a draft genome assembly is available for *C. wangi*, which was generated by Wang et al. (2023) using Oxford Nanopore Technologies (ONT) long-read sequencing, with an estimated genome size of approximately 1.61 Gb. However, a high-quality chromosome-level assembly has not yet been reported for this species. This limitation restricts investigations into the genomic basis of colour variation and comparative genomic analyses within agamid lizards. In the present study, we generated a chromosome-level genome assembly of *C. wangi* using PacBio HiFi long reads and performed Hi-C-based scaffolding. We characterized major genomic features and evaluated assembly quality using multiple approaches. Comparative genomic analyses further identified expanded gene families associated with biological processes and molecular functions potentially related to phenotypic diversification. This high-quality genome assembly provides an important genomic resource for future studies of colour variation and evolutionary diversification in agamid lizards.

## Methods and results

### Sample collection

All animal sampling procedures complied with relevant laws and institutional guidelines for the care and use of animals. Sampling and experimental protocols were approved by the Ethical Committee of China Jiliang University (Approval No. 202560).

A healthy adult female was collected by hand from Changjiang Li Autonomous County (19.2706° N, 109.072° E; 143 m a.s.l.), Hainan Province, China, in November 2024. The individual was anesthetized via intraperitoneal injection of ethyl carbamate (500 mg/kg) prior to tissue collection. Liver tissue was subsequently harvested for genomic DNA extraction. Genomic DNA (gDNA) was extracted using the Yeasen DNA Extraction Kit (Yeasen, Shanghai, China), following the manufacturer’s protocol. DNA quality and integrity were assessed by 1% agarose gel electrophoresis and quantified using a Qubit 4.0 fluorometer (Thermo Fisher Scientific, MA, USA).

An additional adult individual was collected for RNA sequencing in June 2025 from Ledong Li Autonomous County, Hainan, China (18.552° N, 109.095° E; 143 m a.s.l.). Total RNA was extracted from multiple tissues, including heart, liver, gonads, brain and muscle using TRIzol reagent (Invitrogen, Thermo Fisher Scientific, USA) according to the manufacturer’s instructions. RNA integrity was evaluated using 2% agarose gel electrophoresis, and concentration was measured with a Qubit fluorometer (Thermo Fisher Scientific, MA, USA).

### Genome sequencing

To improve assembly continuity and base-level accuracy, we generated PacBio HiFi long reads and performed Hi-C-based scaffolding in the present study. For long-read sequencing, gDNA was used to construct SMRTbell libraries and sequenced on the PacBio Sequel II platform (PacBio, Menlo Park, CA, USA) in Circular Consensus Sequencing (CCS) mode to generate HiFi reads. A total of 98.4 Gb of CCS data (∼59.2× coverage) was obtained, with a mean read length of 14.8 kb. These HiFi reads were assembled into primary contigs using Hifiasm (v0.19.2) with default parameters, resulting in a draft genome assembly of approximately 1.71 Gb with a contig N50 of 110.09 Mb. For transcriptome sequencing, RNA samples were performed on the Illumina NovaSeq 6000 platform using paired-end 150 bp reads, generating a total of 31.18 Gb of raw data. These data were subsequently used to support coding-gene prediction.

### Hi-C chromosome anchoring

Hi-C sequencing was performed to anchor and orient the draft contigs, enabling chromosome-level genome assembly of *C. wangi.* Approximately 1 g of liver tissue was fixed with 1% formaldehyde to preserve chromatin interactions. Genomic DNA was digested with DpnII (a 4-base cutter restriction enzyme), and fragment ends were repaired and labeled with biotin. The labeled DNA fragments were ligated to form chimeric junctions, followed by reversal of crosslinking, DNA purification, and mechanical shearing into 300-500 bp fragments. Biotin-labeled fragments were enriched using streptavidin-coated beads to construct the Hi-C library. The Hi-C library was sequenced on the Illumina platform, generating a total of 139.0 Gb of data. Raw reads were filtered using fastp (v0.23.2) to remove low-quality reads and adapter sequences. The clean reads were processed using the HiC-Pro (v3.1.0) to remove invalid pairs, and valid pairs were used to generate Hi-C interaction matrices using Juicer (v1.6). Contigs were subsequently anchored and oriented into chromosomes using the 3D-DNA (v180922) pipeline, followed by manual curation in Juicebox v1.11.08. In total, the final chromosome-level comprises 6 macrochromosomes and 11 microchromosomes (Fig. 1A and B), totaling 1.66 Gb in length. Furthermore, our assembly achieved a significant improvement in contiguity, with a scaffold N50 of 231.83 Mb, compared to the 91.60 Mb contig N50 of the previous draft^16^.

**Fig 1.**
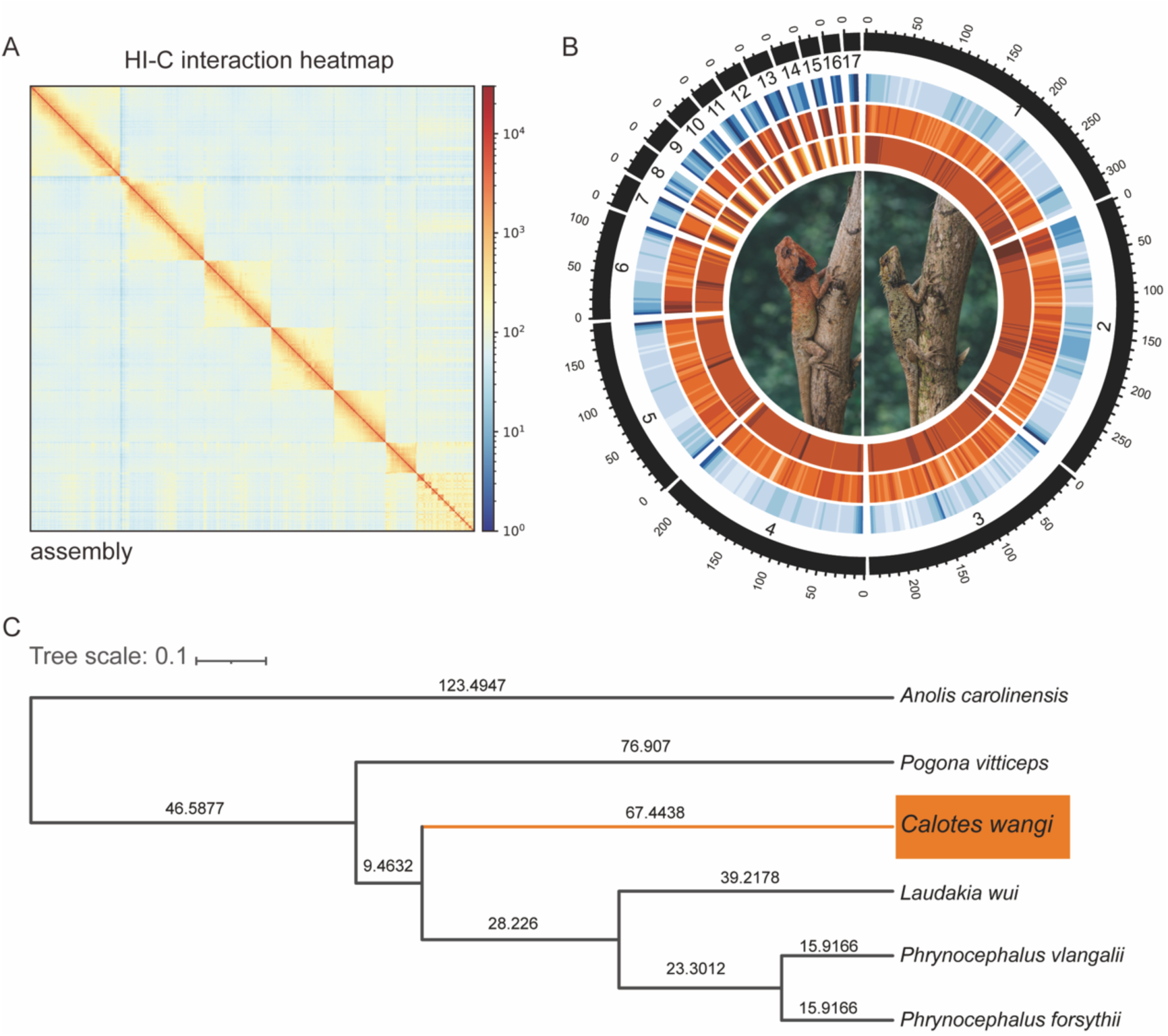
Overview of *Calotes wangi* and its genome features. A. Hi-C interaction heatmap showing chromosome-scale assembly; B. Circular plot showing the gene features. C. Phylogenetic analysis of six lizard species was performed, with *Anolis carolinensis* used as the outgroup.

### Genome annotation

Repetitive elements in the genome were identified using both *de novo* and homology-based approaches. RepeatModeler (v2.0.3) was used to identify interspersed repetitive elements. Long terminal repeat (LTR) retrotransposons were identified by integrating the outputs of LTRharvest v1.6.5^17^ and LTR_FINDER_parallel v1.1^18^. The identified LTR candidates were further refined using LTR_retriever v3.0.1^19^. The resulting repeat libraries were merged and filtered to remove redundant sequences and elements shorter than 50 bp. The curated repeat library was subsequently used for *de novo* repeat annotation with RepeatMasker. Known repetitive elements were annotated using RepeatMasker v4.1.4^20^ against the Dfam v3.6 database^21^ with the Squamata taxonomy. In total, repetitive sequences accounted for approximately 53.39% of the genome, with transposable elements (TEs) representing the predominant repeat category. Among these, long interspersed nuclear elements (LINEs) were the most abundant, comprising 14.48% of the genome.

Protein-coding genes were predicted using BRAKER v3.0.8 with integrated evidence from ab initio prediction, homology-based annotation, and transcriptome-supported evidence. The BRAKER pipeline iteratively trained GeneMark-ETP v1.02^22^ and AUGUSTUS v3.5.0^23^. RNA-seq reads were first aligned to the genome using STAR v2.7.10b^24^. For homology-based annotation, orthologous protein sequences from Tetrapoda were retrieved from OrthoDB v11^25^. Protein sequences from five closely related species, including *Laudakia wui* (GCA_977999105.1^9^), *Pogona vitticeps* (GCA_900067755.1^26^), *Phrynocephalus vlangalii* (GWHBCKV00000000^27^), *P. forsythii* (GCA_029282475.1^28^), *Anolis carolinensis* (GCA_035594765.1^29^), were also incorporated. In addition, all available Squamata protein sequences were downloaded from the UniProt Knowledgebase (UniProtKB, downloaded on July 14, 2025)^30^.

To further annotate untranslated regions (UTRs), the PASA pipeline v2.5.3^31^ was employed. Transcript assemblies generated by Trinity v2.15.2^32^ were aligned to the genome using minimap2 v2.24^33^. PASA was subsequently used to update gene models. A total of 20,442 protein-coding genes were predicted, with an average gene length of 39,341 bp. This represents a substantial improvement over the previous annotation containing 17,547 genes^16^. Genome annotation completeness was assessed using BUSCO (v5.2.2) with the vertebrata_odb10 dataset. A total of 98.9% of BUSCO genes were identified as complete, with 0.6% fragmented and 0.5% missing.

Functional annotation of predicted genes was performed by aligning protein sequences against public databases, including the NCBI Non−Redundant (NR) database and Clusters of Orthologous Groups (COG), using BLASTP with an E-value cutoff of 1e-5. For each sequence, only the best hit was retained to ensure annotation accuracy. Gene Ontology (GO) terms were assigned InterProScan (v5.34-73.0) and direct mapping of Swiss-Prot accessions to GO terms using the IDmapping file. In total, 18535 genes were successfully annotated, and 17,597 Gene Ontology (GO) terms were assigned.

### Comparative analysis of phylogeny and gene families

Phylogenetic relationships and gene family evolution were analyzed using the same five representative reptilian species used for homology-based annotation. Orthologous gene families were identified using OrthoFinder (v2.5.4) with BLASTP (v2.15.0, E-value ≤ 1e-5). Single-copy orthologs genes shared among all species were extracted for downstream analyses. Protein sequences of single-copy orthologs were aligned using MAFFT (v7.490) and trimmed with trimAl (v1.5). The corresponding CDS alignments were subsequently generated and concatenated. A maximum likelihood phylogenetic tree was reconstructed using IQ-TREE2 (v2.3.6) with 1,000 bootstrap replicates. Divergence times were further estimated using MCMCTREE implemented in PAML (v4.10.7). The resulting phylogeny placed *C. wangi* within the agamid lineage and supported its phylogenetic position within the Agamidae family. The inferred topology was consistent with previously reported phylogenetic relationships^9,12,16^. Our analysis estimated the divergence between *C. wangi* and the *Laudakia*/*Phrynocephalus* lineage to be approximately 67.44 million years ago (95% HPD: 44.82–93.63 MYA) (Fig. 1C).

Gene family expansion and contraction were analyzed using CAFE (v5.1.0) under a birth-death model based on the reconstructed species phylogeny. Functional enrichment analysis of expanded gene families was performed using clusterProfiler v4.10.1^34^, with a focus on biological process (BP) and molecular function (MF) categories. A total of 297 gene families were identified as significantly expanded in the *C. wangi* lineage (Table 2). In the BP category, expanded gene families were significantly enriched in terms associated with steroid metabolic processes, epigenetic regulation of gene expression, nucleosome assembly, and actin nucleation (Fig. 2A). Several enriched terms were also related to regulation of viral transcription and DNA integration. In the MF category, enriched terms included protein heterodimerization activity, structural constituents of chromatin, DNA polymerase activity, reverse transcriptase activity, and oxidoreductase activity (Fig. 2B). Terms associated with steroid metabolism, including androsterone dehydrogenase and testosterone 17-beta-dehydrogenase activities, were also enriched. These enriched functions suggest that lineage-specific gene family expansion in *C. wangi* may contribute to regulatory and metabolic processes potentially associated with dynamic colour variation and phenotypic plasticity.

**Fig. 2.**
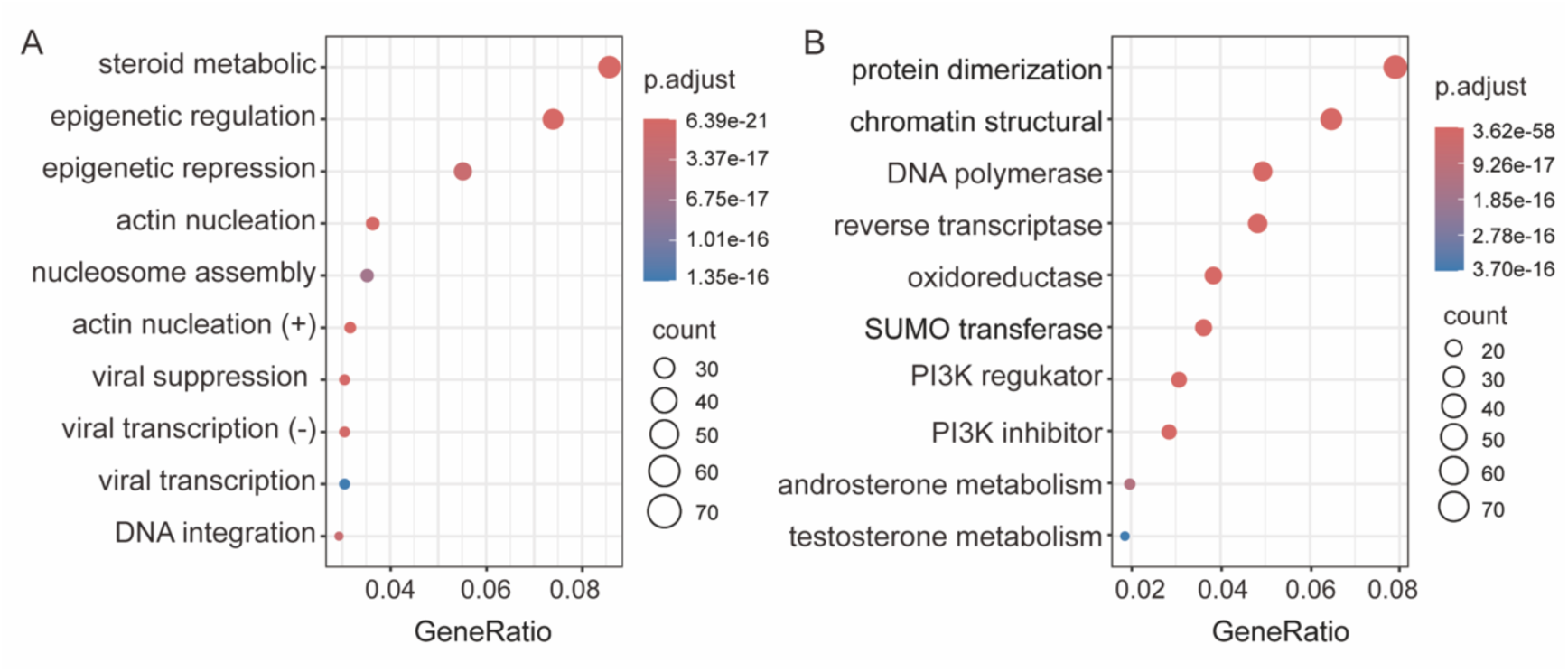
Functional enrichment analysis of expanded gene families in *Calotes wangi.* A. Significantly enriched Gene Ontology (GO) biological process (BP) categories. B. Significantly enriched GO molecular function (MF) categories. Bubble size indicates gene count, and colour represents adjusted p-values.

**Table 1.** Summary of genome assemblies and gene annotations of *Calotes wangi* genome.

| Attribute | Value |
| --- | --- |
| Total length (bp) | 1,662,271,688 |
| N50 (bp) | 231,833,277 |
| GC contents (%) | 43.80% |
| Assembly BUSCO completeness | C:97.9%[S:94.9%,D:3.0%],F:0.6%,M:1.5%,n:3354 |
| Gene number | 20,442 |
| Transcript number | 32,816 |
| Annotation BUSCO completeness | C:98.9%[S:59.1%,D:39.8%],F:0.6%,M:0.5%,n:3354 |

**Table 2.**
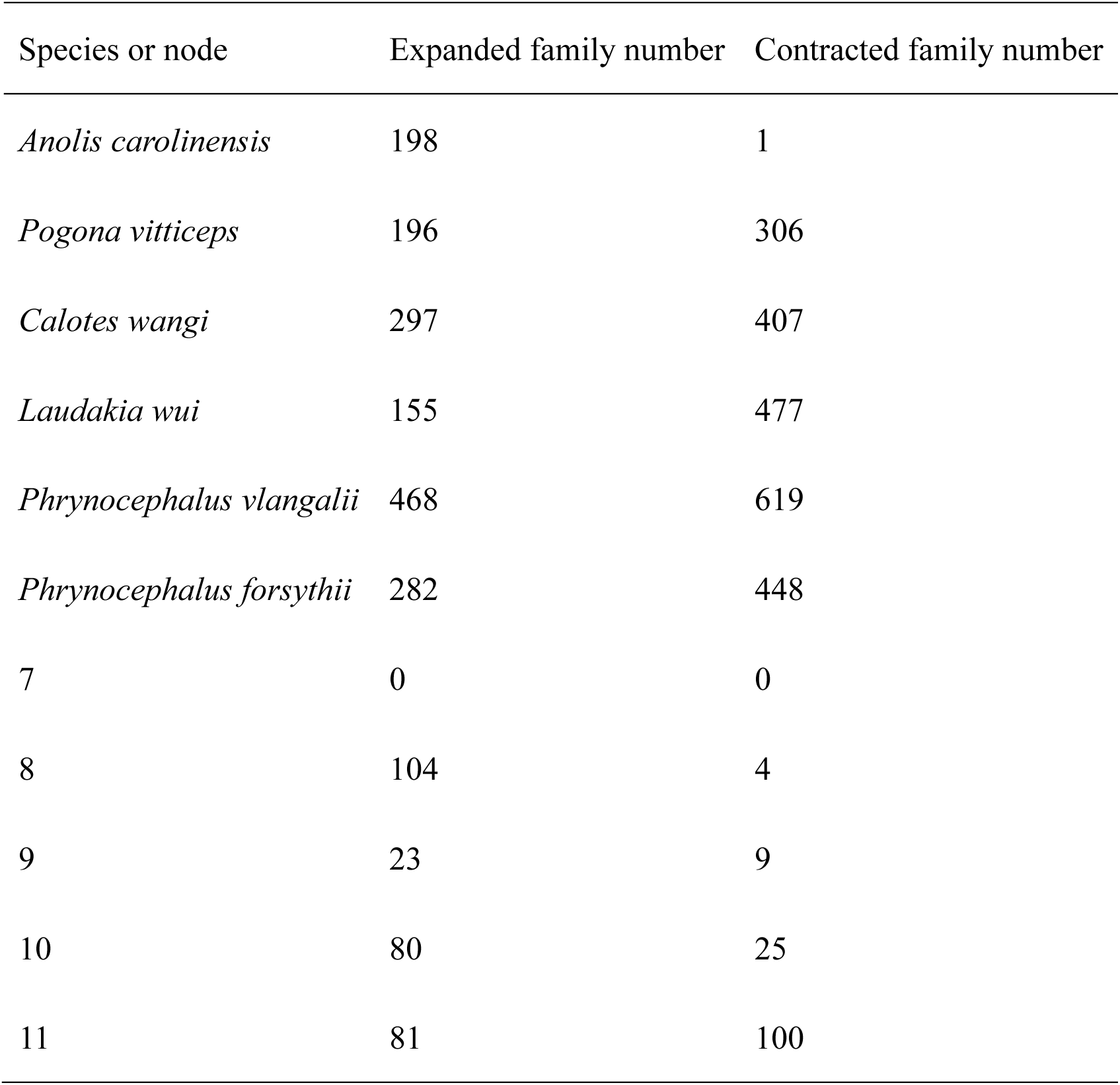
Significantly changed gene families identified in different species.

## Data availability

The data generated in this study are available as follows: the raw sequencing data are available in the NCBI Sequence Read Archive (SRA) under SRP692906; the genome assembly is available at the European Nucleotide Archive (ENA) under JBWZQO000000000; and the annotation files are available from the Figshare repository via 10.6084/m9.figshare.32856524.

## Data Records

The dataset generated in this study includes transcriptome sequencing, PacBio HiFi reads for *de novo* assembly, and Hi-C reads for chromosome scaffolding.

## Technical Validation

Genome assembly completeness was assessed using BUSCO v5.2.2 with the vertebrata_odb10 dataset. A total of 98.9% of BUSCO genes were identified as complete, with 0.6% fragmented and 0.5% missing. BUSCO analysis of predicted protein-coding genes further supported the high completeness of genome annotation. Hi-C interaction heatmaps showed strong interaction signals within each pseudochromosome and clear chromosomal boundaries (Fig. 1A), supporting the accuracy of chromosome anchoring and scaffolding.

## Code availability

All software used in this study is publicly available. Parameters for each analysis are described in the Methods and results section. Unless otherwise stated, default parameters were used.

## Acknowledgements

This work was supported by the National Natural Science Foundation of China (No. 32400381) and Chengdu Institute of Biology, Chinese Academy of Sciences (No. GHZD2025-1). We thank colleagues from Hainan Normal University for assistance with field sample collection. We thank Dr. Yanqing Wu from Wenzhou University for providing species photographs. We are also grateful to Dr. Yanyan Zhou for valuable discussions on genome results. In addition, we thank OpenAI for assistance with language polishing.

## Author contributions

Xia Qiu: conceptualization, data curation, investigation, supervision and writing—original draft, writing—review and editing; Yiru Wang: data curation, formal analysis, investigation, methodology, software, validation, visualization, writing—Original draft; Jieru Wen: data curation, software, visualization; Ying Chen: data curation, software, visualization; Longhui Zhao and Jiayi Jian: revised and edited the manuscript; Weizhao Yang: conceptualization, investigation and writing—original draft, writing—review and editing. All authors read and approved the final manuscript.

